# Turning the screw: engineering extreme pH resistance in *Escherichia coli* through combinatorial synthetic operons

**DOI:** 10.1101/2020.02.12.946095

**Authors:** Guilherme M. V. de Siqueira, Rafael Silva-Rocha, María-Eugenia Guazzaroni

## Abstract

Adoption of microorganisms as platforms for sustainable biobased production requires host cells to be able to withstand harsh industrial conditions, which are usually far from the ones where these organisms are naturally adapted to thrive. However, novel survival mechanisms unearthed by the study of microbiomes from extreme habitats may be exploited to enhance microbial robustness under the strict conditions needed for different applications. In this work, synthetic biology approaches were used to engineer enhanced acidic tolerance in *Escherichia coli* under extreme conditions through the characterization of a library of twenty-seven unique operons composed of combinatorial assemblies of three novel genes from an extreme environment and three synthetic ribosome binding sites. The results here presented illustrate the efficacy of combining different metagenomic genes for tolerance in truly synthetic genetic operons, as expression of these gene clusters increased hundred-fold the survival percentage of cells exposed to an acidic shock in minimal media at pH 1.9 under aerobic conditions.

## Introduction

Biotechnology plays a central role in the expanding search for sustainable solutions to mitigate industries’ dependency on non-renewable substrates, and the production of diverse chemicals by microorganisms has been of great importance in providing alternatives to already established petroleum-based processes. Microorganisms can be exploited as microbial cell factories due to their natural features, or metabolically engineered to produce both chemical building blocks and high added value products from renewable feedstocks such as carbohydrates, glycerol and single carbon compounds ^1^. However, despite the richness of microbes that inhabit the most diverse habitats all over the planet, only a fraction of them can be properly cultivated in laboratory^2^, and the establishment of new industrial microbial processes has been limited to even fewer organisms which possess desirable fermentative characteristics ^3,4^. A major challenge for these endeavors is that microorganisms often lack both the metabolic flexibility and robustness to endure the usually strict and temporally dynamic conditions in parameters such as temperature, pH and even product concentrations that are associated with sustainable and cost-effective industrial fermentations, hindering the replacement of petroleum-based processes by microbial production^5,6^.

Even if not all microorganisms may be cultivated by classical microbiological approaches, technologies that have enabled their genetic material to be accessed and explored have grown exponentially in recent years and have granted researchers novel tools for the study and exploitation of these genomes to diverse ends^7^. Metagenomic studies have been of great importance to uncover proteins with superior industrial properties^8^, rare bioactive compounds^9^, and even novel genetic parts for the construction of reliable synthetic circuits in non-conventional bacterial hosts^10^; moreover, studies from the metagenome of contaminated sites have revealed novel pathways for the degradation of heavy metals^11^, toxic chemicals^12^, as well as innovative mechanisms that enable microorganisms to thrive in the most hostile environments^13^, which can be tapped to grant industrially relevant microorganisms the ability to perform under the harsh conditions needed for efficient fermentation conditions.

The use of one key protein or larger natural protein complexes derived from extremophile microorganisms has already been used to develop promising strains showing improved tolerance levels to different insults, such as temperature, solvent concentration, acidity and oxidative stress^14–16^, and show the feasibility of exploiting these natural mechanisms to engineer robustness in microbial hosts. In this work, we investigate the effectiveness of expressing synthetic assemblies of different acid resistance determinants by tailoring a set of 27 small operons (comprising less than 1.4kb in size each) that consisted of three novel proteins previously uncovered from an extremely acidic environment^17^ regulated by different <u>R</u>ibosome <u>B</u>inding <u>S</u>ites (RBS). The operons were investigated regarding their potential to enhance *Escherichia coli* survival under a pH 1.9 acidic shock, as well as the fitness cost that their expression exerted over the cells. Our results show that, although individually the genes displayed modest capacity to confer acid resistance, the survival percentage of cells was sharply enhanced by their simultaneous expression, and that a wide range of survival phenotypes was achieved by simple permutations of key genetic elements. Yet, the results showed a non-linear relation between protein translation and observed resistance levels, as increasing RBS strength not always resulted in increased acid tolerance. Altogether, the approach presented here allowed the identification of strains with diverse growth profiles that were over 100-fold more tolerant to the acidic stress than the original parental one, demonstrating the potential of metagenomics and synthetic biology to expand the cell capabilities.

## Results and discussion

### Operon library assembly

Using functional metagenomic libraries, we previously identified a collection of different genes from acidic river samples that enabled *E. coli* and other bacteria to withstand an extremely acidic challenge ^17^. From this set, three genes were chosen to compose the synthetic operons engineered in this study. Namely, these genes were annotated as bearing similarities to the DNA-binding protein HU, a major component of nucleoid in bacteria^18^, to a RNA-binding protein (RBP), as well as to the ATP-dependent serinoprotease ClpP, a member of the chaperone-protease complexes that assist misfolded protein degradation as a mechanism of stress response in diverse organisms ^17,19^. In order to assess whether these novel metagenomic genes could work together to further improve bacterial tolerance to acidic stress, a combinatorial approach was employed for the engineering of synthetic operons (Acid Resistance Circuits, ARCs) composed by the three different resistance genes and three synthetic RBS over a pSEVA232 backbone^20^ (**Figure 1; Table 1**).

**Figure 1.**
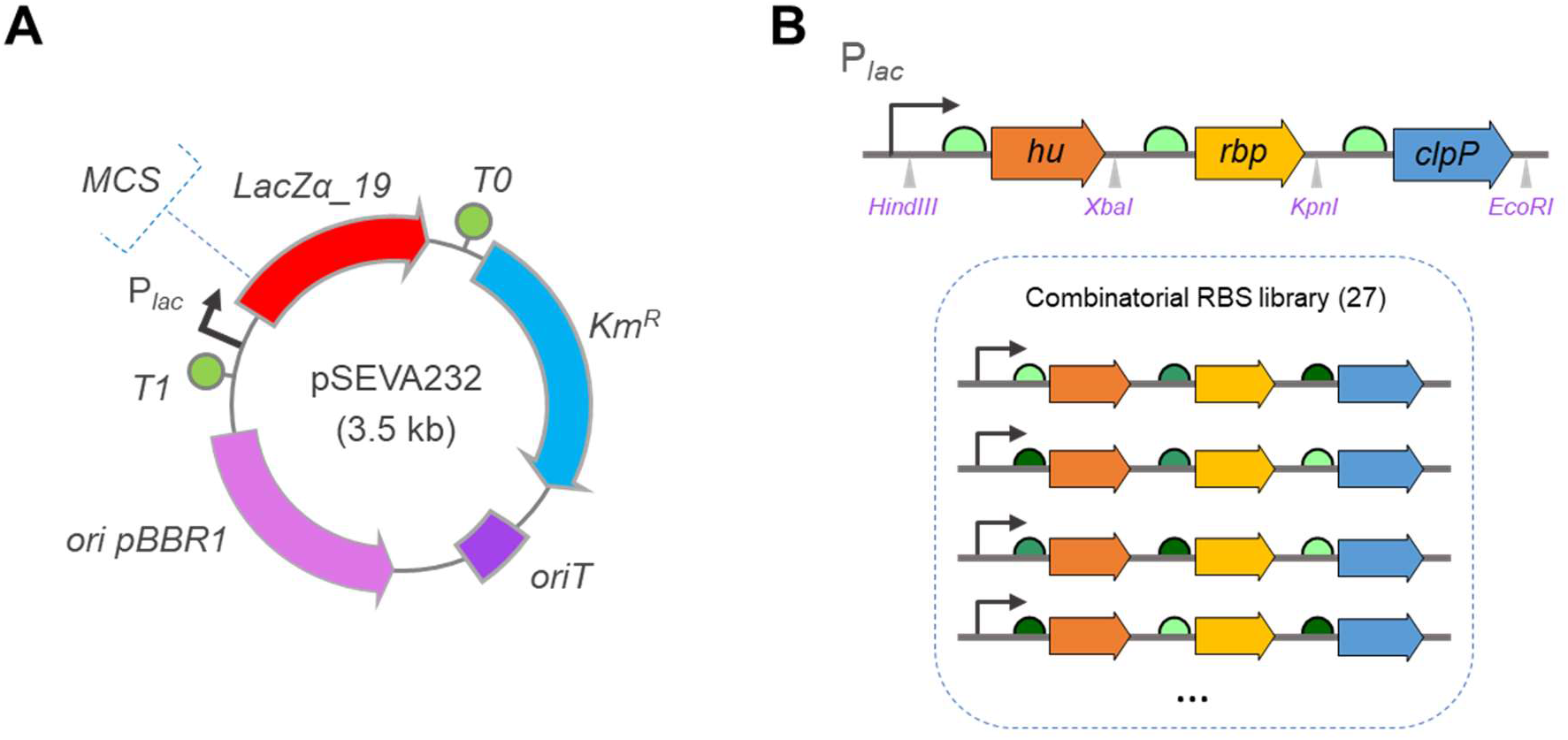
Schematic representation of the strategy employed to build the acid resistance circuits, ARCs. **A)** Circuits were cloned in the multiple cloning site (MCS) of vector pSEVA232^20^. Functional elements of the plasmid backbone are shown: Km^R^. antibiotic resistance marker; *oriT*. origin of transfer; *ori* pBBR1, broad host-range origin of replication; *T1* and *T0*, transcriptional terminators. **B)** Representation of synthetic circuits’ architecture, showing each element in its respective predicted position. In this and every subsequent figure, the curved arrows depict the P_*lac*_ promoter present in the pSEVA232 vector, the colored thick arrows represent the different stress resistance genes, and the semicircles represent the RBS. A scale of green was assigned to the RBS, and color intensity varies to represent their reported strengths of translation: light green is RBS1 (Bba_B0031), medium green is RBS2 (Bba_B0030) and dark green is RBS3 (Bba_B0034). According to this key, the circuits shown in Figure **1B** are pARC111 (top), pARC123, pARC321, pARC231 and pARC313 (bottom). Restriction sites used for cloning are shown in pink.

**Table 1.**
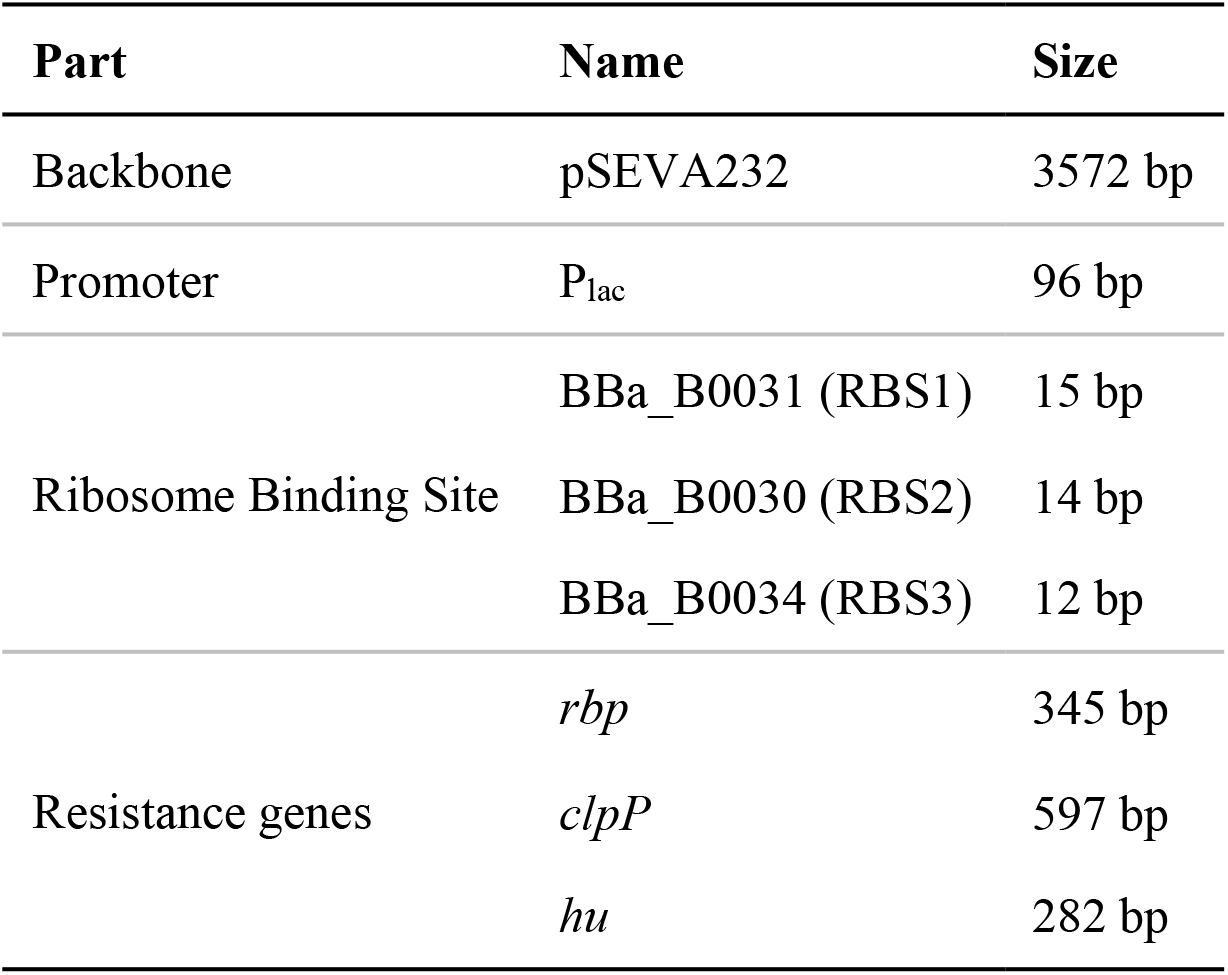
Summary of the biological parts that constitute the genetic circuits built in this work.

In these genetic circuits, the relative gene positions were fixed, *hu* being the first gene in the polycistronic mRNA, followed by *rbp* and then *clpP*, and an RBS slot was available upstream to each of them, allowing for the customization of the operon with any of three selected RBSs as a way to fine-tune translation levels between the different proteins (**Figure 1**). Accordingly, a calibration of the different RBSs using GFPlva, a fluorescent protein possessing a C-terminal degradation tag ^21^, as a proxy for gene expression was performed to determine their suitability to our system and revealed that the RBSs were both functional and, under these circumstances, allowed the distinction of low, medium and high relative expression profiles **(Figure S1).** Ultimately, this combinatorial approach resulted in 27 unique operon designs which were referred to as pARCXYZ, being “X”, “Y” and “Z” indicators of which RBS was present in each slot of the polycistronic mRNA in a given construction. **Figure 1B** provides an illustration for the general design of the synthetic circuits, while **Table 1** presents an overview of all the biological parts used in this work.

### Expression of synthetic operons greatly enhanced survival of bacteria under acidic challenge

The extent to which the expression of *hu*, *clpP* and *rbp* genes was able to confer acid tolerance to *E. coli* was assessed under to strict nutritional and physiological conditions in order to diminish native tolerance responses known to play important roles in *E. coli* stationary-phase survival^22^. For this, each gene was separately expressed under the control of a strong RBS sequence, and exponentially growing cells possessing these constructions were subjected to a 1.9 pH acidic challenge in minimal media (**Methods**). In our assays, we observed that the individual expression of genes granted acid tolerance to the cells in differing degrees, with cells expressing *hu* displaying a superior performance over the ones expressing either of the other genes over, with 0.9% of survival after 1h of acidic challenge. This level was more than 10 times the 0.06% survival of *E. coli* cells harboring only an empty plasmid without tolerance genes. Yet, the survival rate for this clone showed a sharp decay after 2 hours of incubation, as it fell to 0.1% under these conditions (**Figure 2; Table S1)**. Interestingly, when the three genes were co-expressed into an operon under the control of a weaker RBS upstream of each gene (e.g. pARC111), we were able to obtain 9.3% of survival percentages after one hour of acidic shock, resulting into a 10-fold improvement over the expression of *hu* by itself. Additionally, a much smoother decay rate in survival percentages was observed after two hours of acid exposure, as survival fell to 4.10% (**Figure 2; Table S1)**. Such an improvement suggests that the relationship between gene expression and acid tolerance may not only be related to the absolute amount of protein translated, but that a synergic component played an important role in improving bacteria survival under prolonged periods of stress.

**Figure 2.**
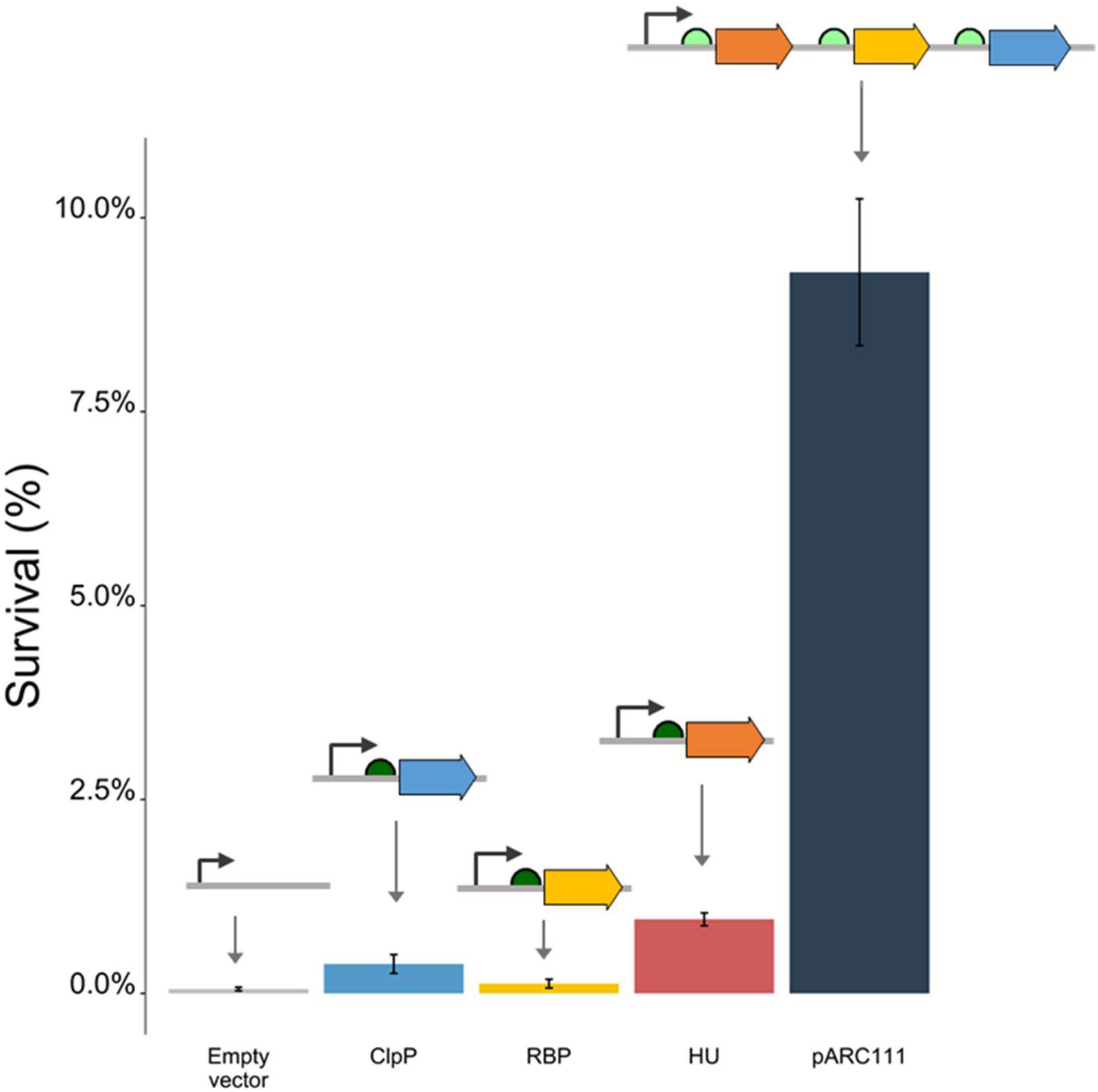
Acid resistance of the clones harboring plasmids with the individual resistance genes *clpP, rbp* and *hu* under the control of RBS3 and with pARC111, the operon which contains the weakest combination of RBS, showcasing the advantage of concurrent expression of such genes. Schematic representations of the synthetic circuits are shown above their respective bars. The survival is calculated as the percentage of colony forming unities after 1 h exposure to pH 1.9. Error bars indicate standard deviation from three independent experiments.

Next, we investigated how further combination of the genetic parts impacted the observed survival levels. As seen in **Figure 3**, if pARC111 is to be considered a ‘basal activity’ operon, the change of relative protein levels due to the replacement of the RBS in each position led to different and unexpected survival profiles. For instance, the exchange of RBS1 to RBS2 in the position of *clpP* in pARC112 caused no absolute difference in bacterial survival when compared to pARC111, while the presence of RBS3 in the same position more than doubled the observed survival rates (as seen in pARC113). This relationship is not always true, however, since the RBS1 to RBS2 replacement in the *hu* position had a more significant impact than RBS1 to RBS3, as the pARC211 expressing strain showed almost 50% of survival after one hour of incubation whereas pARC311 cells survival remains at a 30% level. Given that *hu* was also the individual gene associated to the greater levels of survival, these results may reinforce that genes play different roles in enhancing cells’ resilience under stress, but they also hint to the context-dependent nature of RBS activity, as the tridimensional architecture derived from the combination of the RBS sequence itself and its surrounding regions (genetic context) has been shown to influence the recruitment and movement of ribosomes through the mRNA^23^. Furthermore, considering that our constructs are all expressed as a polycistronic mRNA strand, it must also be considered that the translation rates of the genes are not entirely independent, and relative RBS strength may vary as a result of the translational coupling between the adjacent genes^25^, a phenomenon that adds another whole layer of complexity to the 27 combinations that were made.

**Figure 3.**
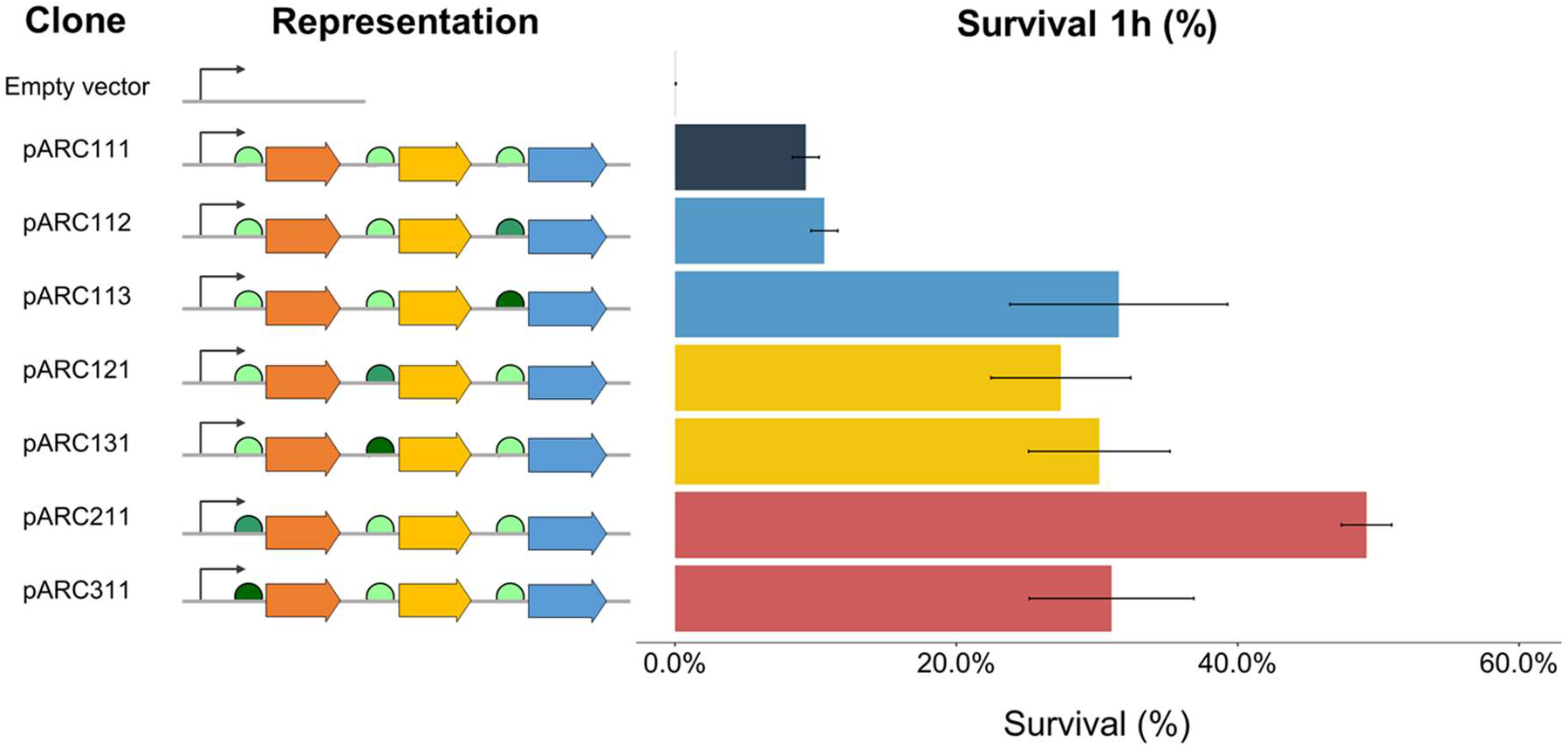
Differences in acid resistance promoted by variation of a single RBS in the synthetic circuits. Dark blue bar shows the survival percentage of pARC111, a “neutral” circuit, after 1h under acidic challenge; light blue bars, yellow bars and red bars, respectively, depict survival percentages of clones in which the RBS of *clpP*, *rbp* and *hu* were altered. Error bars indicate standard deviation from three independent experiments.

### Survival percentages and expression costs for host cells varied due to operon design

A wide range of survival profiles was obtained through the expression of our group of synthetic operons, as shown in **Figure 4**. Interestingly, cells bearing constructions one could argue as possessing the strongest set of tolerance determinants (i.e. strong RBS in every gene, such as pARC333, pARC323, etc.) showed lower survival percentages than those carrying more evenly composed operons, which we hypothesize to be a consequence of a heavier expression burden imposed over these cells than over those expressing less-demanding constructions^26^. Additionally, analysis of cellular growth profiles showed that the growth of strains bearing any of the plasmids containing pARCs was impaired when compared to the growth of a control carrying an empty plasmid, even at lower pH, despite the expression of pARCs evidently enabling great improvement in survival under acidic stress **(Figure S2; Table S2)**.

**Figure 4.**
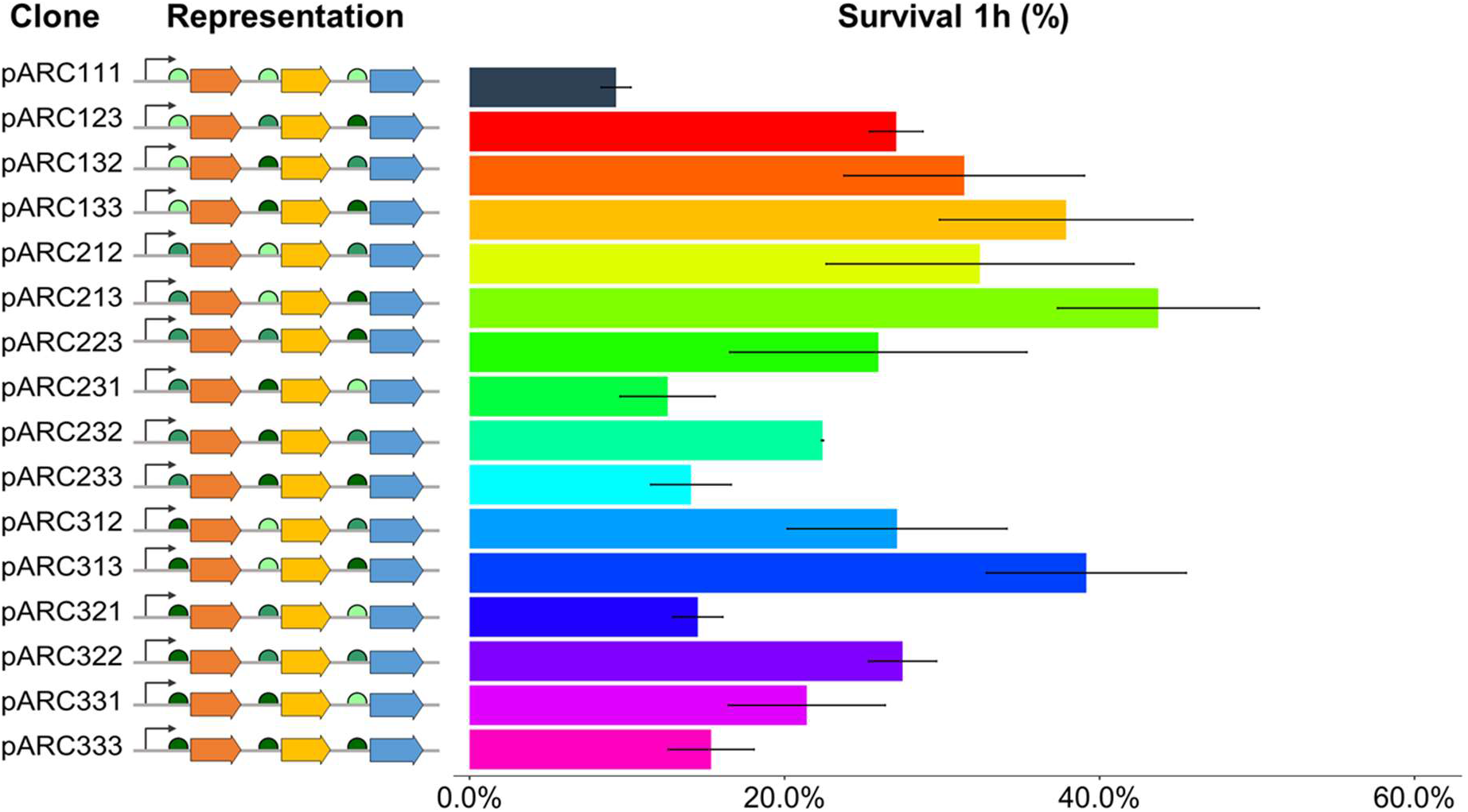
The combination of resistance genes under translational control of different synthetic RBS allowed that wide range of acid resistance levels could be obtained, as shown by the survival percentages calculated after the acidic challenge of clones harboring pARCs. Error bars indicate standard deviation from three independent experiments.

Aside from the quantitative burden of expressing any number of exogenous proteins, this behavior might suggest that the mechanism through which pARCs promote stress tolerance may involve a trade-off with cells’ multiplication capacity, which may be consistent to putative stationary-phase roles that the expressed proteins may play. For instance, HU is described as a DNA-binding protein that shows affinity towards aberrant DNA formations rather than specific sequences^27^, and has been proposed to have protective effects against DNA damage in radioresistant bacterial species by tightly binding the genetical material together and avoiding the dispersion of fragments after irradiation, allowing efficient repair by the cells^28^. However *E. coli*’s native HU has also been associated with a number of important changes in the transcriptional profile of cells under different stresses and has been shown to positively influence translation of the stationary-phase sigma factor RpoS^18,29^, it is unclear, however, if the expression of this novel HU provides *E. coli* cells with the same capacities as the native one, since *E. coli* and other enterobacteria possess unique heterodimeric HU, as the transcription factor IHF is, whereas HU homologs of other bacteria are usually homodimeric^27^. Given the cost of expressing the heterologous proteins and the interference they may cause in native cellular metabolism, we hypothesize that the prolonged cultivation periods needed to harvest cells prior to the acidic shock assay could have had a selective role in cultures expressing metabolically-demanding operons, and this might have sacrificed the tolerant phenotype by biasing the community towards faster growing cells^30,31^, resulting in underperforming populations, which might explain the observed results in clones bearing constructions such as pARC231, pARC233 and pARC333.

Although this apparent trade-off between resistance and growth was observed, it is remarkable that the comparative analysis of relative fitness between cells harboring pARCs reveals a rich, non-linear relationship between acid tolerance of cells and growth performance, as measured by the decrease in growth rate promoted by the expression of the synthetic operons. This two-dimensional analysis allowed us to explore an expression space and distinguish, between circuits that conferred similar levels of acidic resistance under stress, and those that have achieved this feat with the lowest detrimental effects to the host cells. In this sense, **Figure 5** and **Figure S3** show the relationship between the fitness cost and survival for some of the strains engineered in this work at different time points. For instance, the comparison of pARC111, pARC112 and pARC211 profiles show that, despite showing no difference in survival percentage after the acid resistance assay, the expression of pARC112 was much more costly to the host’s metabolism than pARC111, and the expression of pARC211 provided much more tolerance than pARC112 without a corresponding increase in fitness cost to the cells. Moreover, these results imply that greater levels of survival might be related to the expression of low-cost, efficient operons, as cells expressing pARC211, pARC212 and pARC213 operons, despite displaying an astounding increase in survival in the acid shock assays, showed relatively modest fitness cost. Taken together, this analysis shows how synthetic operons can be exploited to identify optimal solutions to improve bacterial resistance to stress without compromising cellular fitness.

**Figure 5.**
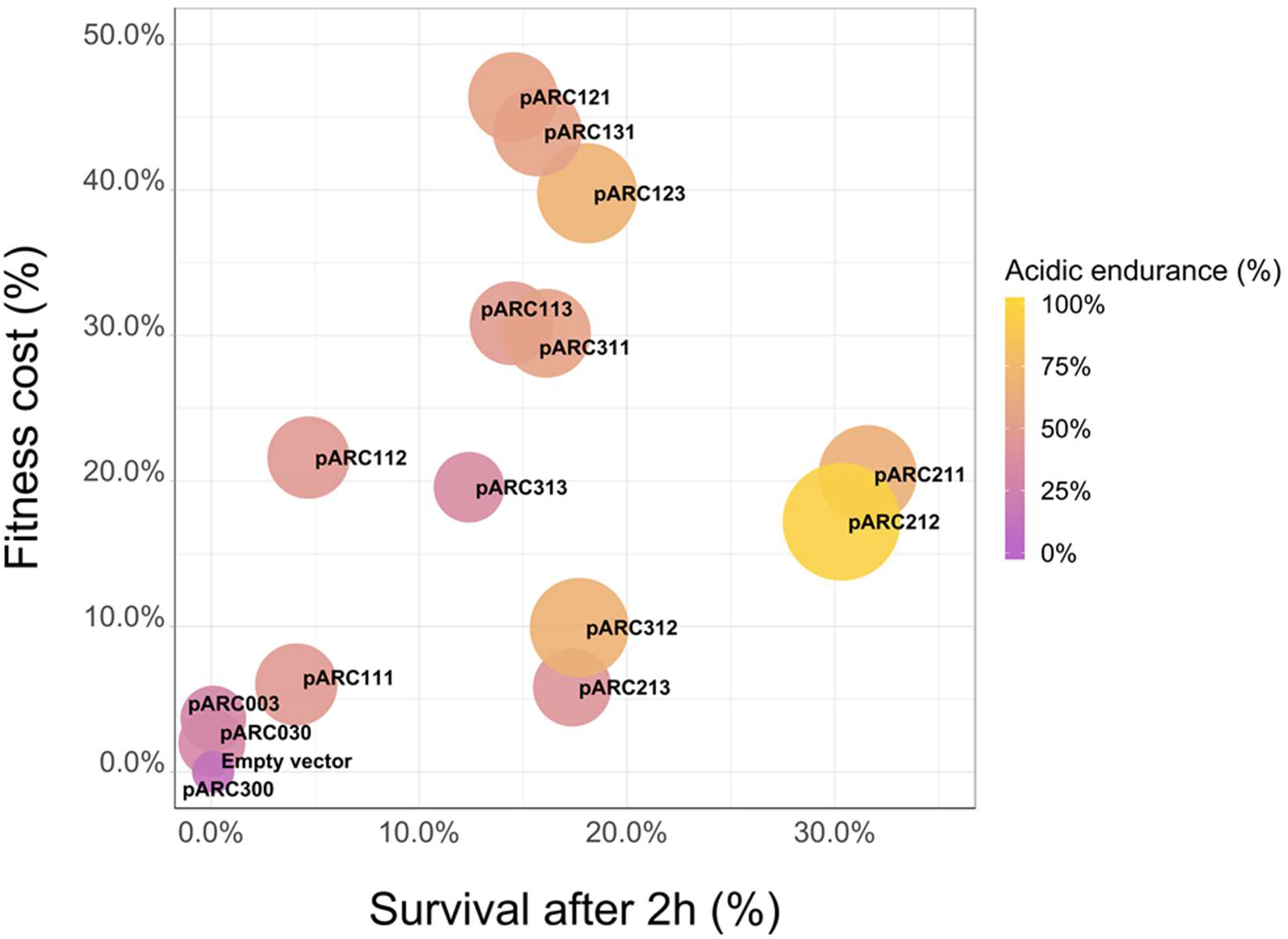
Analysis of the relationship between acid resistance after 2h of acidic challenge and fitness cost of different clones harboring plasmids with pARCs, showing that similar acid resistance levels can be obtained with varied associated expression costs for host cells.

## Conclusion

Several studies have employed combinatory libraries of regulatory elements for optimizing the heterologous expression of biosynthetic pathways^32–34^, and different molecular strategies haven been used for the rational improvement of bacterial robustness under different conditions, such as temperature, pH and ethanol concentration ^35–38^. However, to the best of our knowledge, this is the first time such a combinatory approach was taken to generate a library of truly synthetic tolerance clusters composed by seemingly unrelated resistance genes. This strategy allowed us to navigate through an expression space of strains with different resistance levels and growth profiles and identify cells with maximized acidic resistance despite low fitness cost. It is worth noting that, as a proof of concept, we only investigated operon design properties at translation levels, but the results here presented still may be further expanded by engineering systems also regulated at transcription level. Feedback-controlled promoters that adjust expression levels to match the insult sensed by the cells, for instance, were proven to be extremely effective in providing cells with enhanced fermentative properties due to lower associated expression burden^39,40^. As Synthetic Biology consolidates its position as a discipline that aims to engineer and expand the limits of life, the appropriation of unique mechanisms uncovered from the known bounds where life is found is of great importance to accelerate the development of new, robust synthetic circuits for addressing industrial and societal needs. Approaches such as the one presented in this study might be of great importance to uncover novel functions and non-obvious synergistic relationships between promising proteins, and hopefully researchers will be encouraged to delve more into the ever-growing richness of metagenomes available in genetic databases.

## Methods

### Bacterial strains and culture conditions

*E. coli* strain DH10B was used as a host in molecular cloning and acid challenge steps. All cultivations were performed under aerobic conditions in either LB or M9 minimal media^41^ supplemented with 0.1 mM casamino acids and 1% (v/v) glycerol as carbon source (M9-gly), at 37°C and 220 rpm. When needed, media acidity was adjusted to the desired pH with 1M HCl and filtered with 0.2μM sterile filters. Selection of pSEVA232 vector was performed by the addition of 50 μg/mL of kanamycin in the culture. IPTG was not added to the media, unless otherwise stated, as we noticed that expression levels above *P*_*lac*_ basal rate impaired cell growth for pARC-expressing strains and were not needed for discernible levels of acidic tolerance.

### Operon construction and cloning

The broad-host range, medium copy number plasmid pSEVA232 was used as the backbone for the assembly and expression of synthetic operons^20^. The previously described^17^ genes were wholly synthetized by Integrated DNA Technologies (IDT) from the deposited GenBank sequences (accession numbers JX219763, JX219770, JX219767, for *rbp*, *clpP* and *hu*, respectively). The RBS sequences were retrieved from the iGEM Community Collection (http://parts.igem.org/) and incorporated on 5’ primers for each gene, allowing the amplification of each of them with the desired RBS sequence. Primers were also designed with terminal restriction sites to direct fragment ligation. **Table S3** contains primer sequences used in this study. For the simultaneous ligation of the genes in the linearized vector, an equimolar pool of the fragments was treated as a single insert in ligation reactions and these were carried overnight at 16°C. Ligation mixtures were transformed in electrocompetent *E. coli* DH10B, which were incubated overnight for the growth of recombinant colonies.

### Acid shock assay

Stress tolerance promoted by the expression of tolerance genes and synthetic operons was pursued for exponential-phase cells grown aerobically in nutrient poor media, as *E. coli*’s native resistance mechanisms to extreme acidity aren’t wired to respond well under these conditions^22^. Cells from a single colony were first acclimated to minimal media in a cultivation in 5mL of M9 media for a day at 37°C and 220 rpm. After that, 5μL of culture were diluted in 5mL of fresh media and let grown overnight (16h) under the same parameters in order to harvest cells. The next day, a 1:50 dilution of the culture was done in fresh media and cells were let grow until reaching mid-log phase (OD_600_ between 0.6 and 0.9 depending on the strain), when the acid assay would start by the dilution of 10μL of the culture in 990μL of acid M9 media (pH 1.9), without antibiotics, in a microcentrifuge tube that was incubated at 37°C e 220 rpm.

At the moment cells were incubated (T_o_), as well as hourly during the acid incubation, 10μL of cells were retrieved for colony forming units (CFU) counting and viable cell estimation, as previously described^17^. The acid shock was stopped by serial dilutions in neutral phosphate-buffered saline (PBS) buffer (pH 7.2) and three 25μL droplets (technical replicates) for each dilution at every timepoint were placed onto M9+agar plates and let grow overnight for colony formation. Survival percentage of cells was calculated by the ratio between CFU ml^−1^ at a given point and the t_o_. Experiments were performed with three biological replicates.

### Growth rates measurement and RBS calibration

Cells’ growth was measured at 600nm in the 96-well plate reader Victor X3 (PerkinElmer, Inc.). In order to do so, cells from a single colony were harvested as described for the acid resistance assay, but the overnight grown culture was instead aliquoted to a 1.0 OD as read by the Biophotometer D30 tabletop spectrophotometer (Eppendorf, Inc.). These aliquots were diluted to 1:10 ratio with either neutral or acidified M9 media and antibiotic in the 96-well plate for a final volume of 200μL. The plate was incubated at 37°C for up to eight hours, and punctual measurements of OD600 were automatically performed every 30 minutes. Every experiment was performed with three biological replicates, and three technical replicates were made for each.

RBS calibration experiments were carried in neutral media as described above, however, for these curves, both optical density at 600 nm (OD) and fluorescence (excitation 488 nm and emission 535 nm) were measured, allowing the determination of fluorescence as a function of cell growth over time and IPTG (100μM) was added to the media in order to enhance fluorescence signal.

### Fitness cost calculation

After growth curves were obtained, growth rates (μ) determination was done by estimating the slope of the curves during the linear exponential growth phase ^42^, and the cost of different constructions over the host cells was calculated by the methodology described by Bienick and collaborators ^43^. Fitness was calculated as the ratio between the reference growth rate (μ_ref_), corresponding to the control expressing an empty plasmid, and the growth of each clone. This value was adjusted so that a theoretical quotient of 1 meant 0% of fitness cost, as shown in **Equation 1**

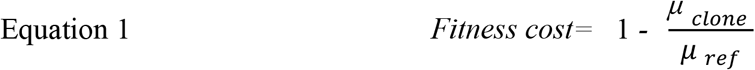

### Data analysis and visualization

Victor X3 data analysis and growth curve elaboration, as well as survival percentages estimations from the acid resistance assay, were made by *ad hoc* scripts using R programming language (version 3.5.2) and the ggplot2 package.

## Author contribution

MEG and RSR conceived and designed the study. GMVS performed the experiments. GMVS wrote the manuscript. MEG and RSR revised the final version of the manuscript. All authors read and approved the final version.

## Supporting information

**Figure S1.** Strength characterization of the synthetic RBS used in this study, as measured by GFPlva fluorescence. **Figure S2.** Growth comparison between clones harboring pARC211 and pSEVA232 empty vector. **Figure S3.** Relationship analysis between acid resistance after 1h and metabolic fitness of the different clones harboring plasmids with combinations of the three RBS. **Table S1.** Survival percentages obtained at time points 1h and 2h of the acidic challenge. **Table S2.** Growth rate and fitness cost of strains grown in minimal medium. **Table S3.** Oligonucleotides used for assembly of synthetic circuits.

## Acknowledgements

The authors wish to thank Thalita Ruil Prado for the invaluable support provided to the realization of our experiments, Caroline Moncaio Moda for assisting the measurements of RBS activity, and to all our lab colleagues for their insightful comments and suggestions throughout the course of this study. This work was supported by the Young Research Awards by the São Paulo State Foundation (FAPESP, award numbers 2015/04309-1 and 2012/21922-8) and was also financed in part by the Coordenação de Aperfeiçoamento de Pessoal de Nível Superior - Brasil (CAPES) - Finance Code 001. GMVS is recipient of a M.Sc. fellowship from FAPESP (award number 2018/07261-8).

