## Supplementary information for "Turning the screw: engineering extreme pH resistance in *Escherichia coli* through combinatorial synthetic operons"

**by**

Guilherme M. V. de Siqueira[1], Rafael Silva-Rocha[2] and María-Eugenia Guazzaroni[3*]

[1] Departamento de Bioquímica, Faculdade de Medicina de Ribeirão Preto (FMRP-USP), Ribeirão Preto, SP, Brasil. 14049-900

[2] Departamento de Biologia Celular e Molecular e Bioagentes Patogênicos, Faculdade de Medicina de Ribeirão Preto (FMRP-USP), Ribeirão Preto, SP, Brasil. 14049-900

[3] Departamento de Biologia, Faculdade de Filosofia Ciências e Letras de Ribeirão Preto (FFCLRP-USP), Ribeirão Preto, SP, Brasil. 14040-901

### Supporting Information

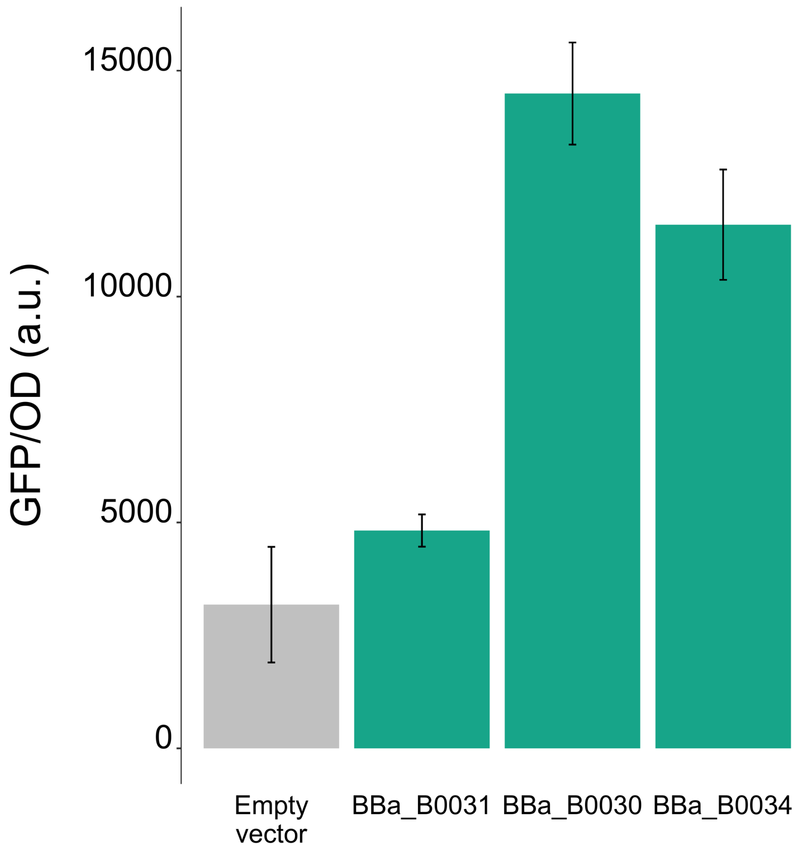

**Figure S1.** Strength characterization of the synthetic RBS used in this study, as measured by GFPlva fluorescence. Considering RBS3 (Bba_B0034) as the standard for 100% translational strength, we calculate relative strengths of 125% for RBS2 (Bba_B0030) and 41,6% for RBS1 (Bba_B0031), which are different values from the ones previously reported in the iGEM database (60% and 7%, respectively). RBS efficiency is heavily dependent on the broader context of mRNA secondary structure, thus the difference between the reported values and the ones obtained in this study may be due to the different genetic contexts in which these RBS are inserted and the conditions in which they were characterized, revealing a non-orthogonal behavior of these genetic parts. a.u.= arbitrary units.

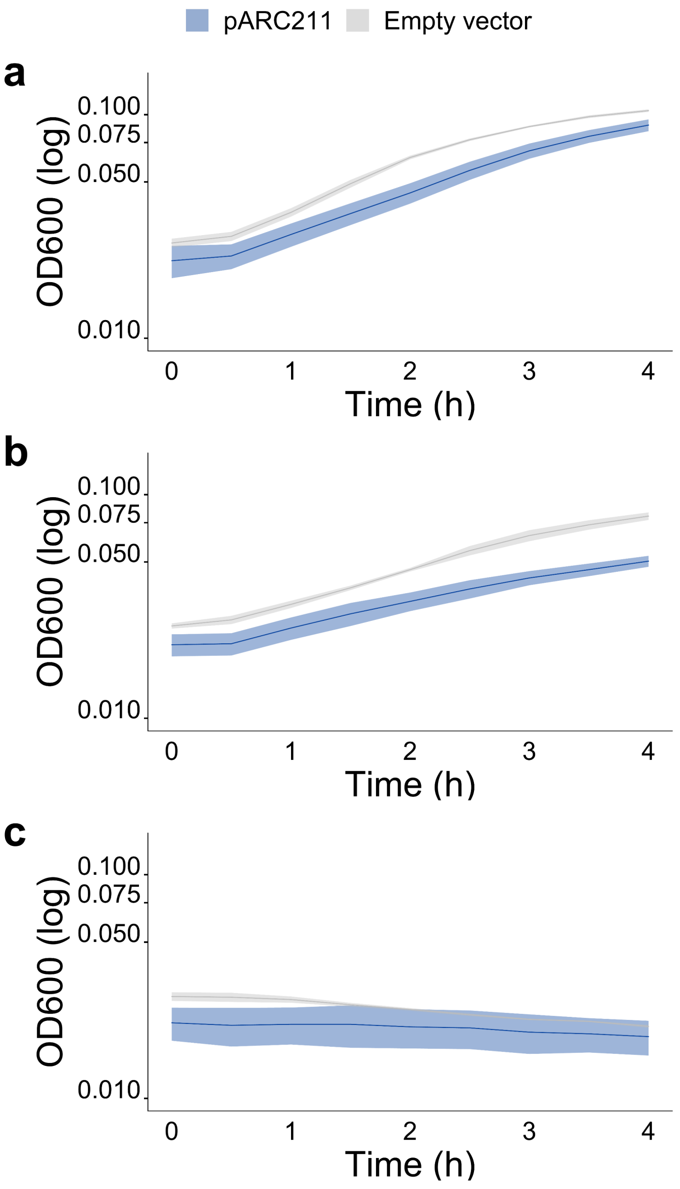

**Figure S2.** Growth comparison between clones harboring pARC211 and pSEVA232 empty vector at **a)** neutral pH; **b)** pH 3.5; **c)** pH 3.

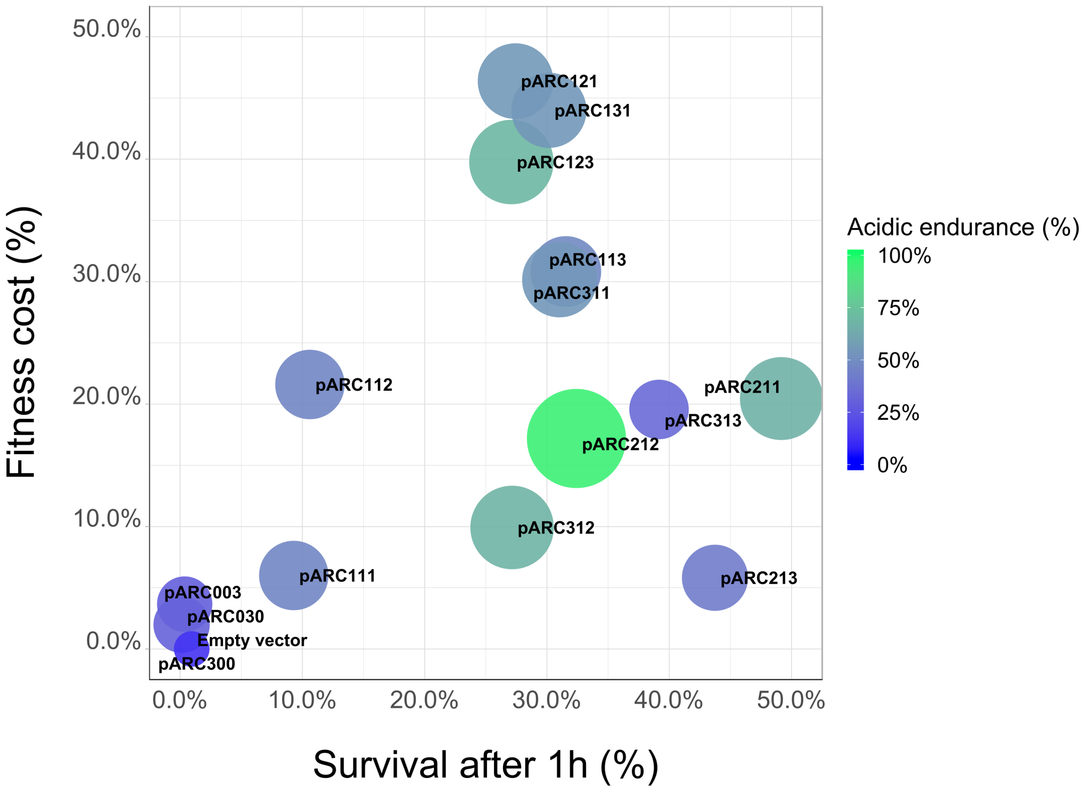

**Figure S3.** Relationship analysis between acid resistance after 1h and metabolic fitness of the different clones harboring plasmids with combinations of the three RBS.

**Table** S1**.** Survival percentages obtained at time points 1h and 2h of the acidic challenge. SP = Survival Percentage, sd = standard deviation

| Strain | SP (1h) | sd (1h) | SP (2h) | sd (2h) |
|---|---|---|---|---|
| pARC000 | 0,06% | 0,02% | 0,00% | 0,00% |
| pARC030 | 0,13% | 0,06% | 0,04% | 0,03% |
| pARC003 | 0,38% | 0,12% | 0,10% | 0,06% |
| pARC300 | 0,96% | 0,08% | 0,10% | 0,03% |
| pARC111 | 9,30% | 0,95% | 4,10% | 0,77% |
| pARC112 | 10,62% | 0,96% | 4,68% | 0,25% |
| pARC231 | 12,57% | 3,01% | 1,88% | 1,87% |
| pARC233 | 14,06% | 2,55% | 7,97% | 2,02% |
| pARC321 | 14,49% | 1,58% | 13,32% | 1,00% |
| pARC333 | 15,33% | 2,73% | 11,14% | 3,93% |
| pARC331 | 21,41% | 4,98% | 17,52% | 0,02% |
| pARC232 | 22,41% | 0,06% | 17,67% | 2,11% |
| pARC223 | 25,96% | 9,43% | 13,37% | 2,31% |
| pARC123 | 27,09% | 1,69% | 18,09% | 1,34% |
| pARC312 | 27,15% | 6,98% | 17,71% | 5,96% |
| pARC121 | 27,43% | 4,97% | 14,52% | 0,23% |
| pARC322 | 27,49% | 2,16% | 15,15% | 1,63% |
| pARC311 | 31,03% | 5,85% | 16,13% | 3,69% |
| pARC132 | 31,41% | 7,63% | 17,30% | 4,28% |
| pARC113 | 31,55% | 7,73% | 14,45% | 3,91% |

| | | | | |
|---|---|---|---|---|
| pARC212 | 32,41% | 9,77% | 30,33% | 9,32% |
| pARC133 | 37,87% | 8,04% | 33,16% | 3,42% |
| pARC313 | 39,16% | 6,34% | 12,41% | 1,18% |
| pARC213 | 43,73% | 6,40% | 17,37% | 0,07% |
| pARC211 | 49,16% | 1,79% | 31,58% | 6,55% |

**Table S2.** Growth rate and fitness cost of strains grown in minimal medium. As stated in the text. growth rates were measured on the linear phase of growth under pH 7 and fitness cost was calculated by the comparison of such rates with the growth rate of the strain used as negative control, which lacks the expression of heterologous genes.

| Strain | Growth rate ($h^{-1}$) | $R^2$ | Fitness cost |
|---|---|---|---|
| pSEVA232 | 0.2732 | 0.9984 | 0.0% |
| pARC003 | 0.2632 | 0.9848 | 3.7% |
| pARC030 | 0.2678 | 0.9931 | 2.0% |
| pARC300 | 0.2743 | 0.9880 | 0.0% |
| pARC111 | 0.2568 | 0.9969 | 6.0% |
| pARC112 | 0.2142 | 0.9904 | 21.6% |
| pARC113 | 0.1890 | 0.9967 | 30.8% |
| pARC121 | 0.1465 | 0.9921 | 46.4% |
| pARC123 | 0.1645 | 0.9938 | 39.8% |
| pARC131 | 0.1530 | 0.9953 | 44.0% |
| pARC211 | 0.2173 | 0.9997 | 20.5% |
| pARC212 | 0.2262 | 0.9948 | 17.2% |
| pARC213 | 0.2573 | 0.9987 | 5.8% |
| pARC311 | 0.1908 | 0.9972 | 30.1% |
| pARC312 | 0.2461 | 0.9999 | 9.9% |
| pARC313 | 0.2198 | 0.9994 | 19.6% |

**Table S3.** Oligonucleotides used for assembly of synthetic circuits. RBS sequences are shown in bold and the annealing sequence to the target gene are underlined. R.S. = Restriction Site

| ID | Sequence (5’ → 3’) | R.S. |
|---|---|---|
| HUF60 | CGCGAAGCTT**ATTAAAGAGGAGAAA**TACTAGATGAGCAAAGAAAAAGAAGTGCT | HindIII |
| HUF7 | CGCGAAGCTT**TCACACAGGAAACC**TACTAGATGAGCAAAGAAAAAGAAGTGCT | HindIII |
| HURev | CGCGTCTAGATTACGCGGCAGTTTTGC | XbaI |
| RBPF60 | CGCGTCTAGA**ATTAAAGAGGAGAAA**TACTAGATGAACAATAAGCTGTACGTTGG | XbaI |
| RBPF7 | CGCGTCTAGA**TCACACAGGAAACC**TACTAGATGAACAATAAGCTGTACGTTGG | XbaI |
| RBPRev | CGCGGGTACCTTAGAAGTGACCGCGACC | KpnI |
| ClpPF60 | CGCGGGTACC**ATTAAAGAGGAGAAA**TACTAGATGGGACTTATACCGATGGTG | KpnI |
| ClpPF7 | CGCGGGTACC**TCACACAGGAAACC**TACTAGATGGGACTTATACCGATGGTG | KpnI |
| ClpPRev | CGCGGAATTCTCAGGTGCGCGGAC | EcoRI |
